# QTL-guided metabolic engineering of a complex trait

**DOI:** 10.1101/079764

**Authors:** Matthew J. Maurer, Lawrence Sutardja, Dominic Pinel, Stefan Bauer, Amanda L. Muehlbauer, Tyler D. Ames, Jeffrey M. Skerker, Adam P. Arkin

## Abstract

Engineering complex phenotypes for industrial and synthetic biology applications is difficult and often confounds rational design. Bioethanol production from lignocellulosic feedstocks is a complex trait that requires multiple host systems to utilize, detoxify, and metabolize a mixture of sugars and inhibitors present in plant hydrolysates. Here, we demonstrate an integrated approach to discovering and optimizing host factors that impact fitness of *Saccharomyces cerevisiae* during fermentation of a *Miscanthus x giganteus* plant hydrolysate. We first used high-resolution Quantitative Trait Loci (QTL) mapping and systematic Bulk Reciprocal Hemizygosity analysis (bRHA) to discover 17 loci that differentiate hydrolysate tolerance between an industrially related (JAY291) and a laboratory (S288C) strain. We then used this data to identify a subset of favorable allelic loci that were most amenable for strain engineering. Guided by this “genetic blueprint”, and using a dual-guide Cas9-based method to efficiently perform multi-kilobase locus replacements, we engineered an S288C strain with superior hydrolysate tolerance than JAY291. Our methods should be generalizable to engineering any complex trait in *S. cerevisiae*, as well as other organisms.

## Introduction

Host development for industrial and synthetic biology applications typically includes breeding, evolutionary engineering, and genetic engineering. These approaches take advantage of pre-existing genetic diversity, generate novel genetic diversity, and/or make use of exogenous genes to construct synthetic pathways^1,2^. Complications in strain development often arise from multiple genetic loci requiring engineering, which frequently exist in complex regulatory, signaling, and/or interaction networks, characteristic of polygenic, complex traits^2–4^. Identifying these targets and engineering them to achieve desired effects while avoiding unintended, off-pathway consequences is challenging, making rational strain design difficult^1–4^.

Industrial production strains encounter a wide variety of natural and non-natural stresses depending on the particular industrial process^1,2,5,6^; therefore, understanding how to engineer complex fitness-based traits, such as stress tolerance, is widely applicable to many bioprocesses. As a test bed for developing complex-trait engineering strategies, we focused on improving *Saccharomyces cerevisiae* strains for bioethanol production from lignocellulosic hydrolysate, which is a complex mixture of fermentable sugars and chemical compounds^7–11^. Some of these compounds act as growth and fermentation inhibitors that prevent the efficient utilization and conversion of sugar into ethanol^7,8,11,12^. Thus, engineering strains with increased tolerance to these inhibitors is an important step towards commercialization of next-generation ethanol and advanced biofuels. We aimed to engineer strains with improved anaerobic growth (a condition that mimics a large-scale industrial fermentation) in the presence of dilute-acid pretreated *Miscanthus x giganteus* hydrolysate (DAH). This hydrolysate was prepared using a harsh pretreatment protocol, and thus contained a high concentration of inhibitors, making it highly toxic^13,14^.

Quantitative Trait Loci (QTL) analysis of an inbred cross is an ideal approach for mapping the genetic basis of complex traits^15^. QTLs can be mapped at high resolution using high-throughput, whole-genome sequencing of pooled individuals from extremely large inbred populations^16,17^. Using QTL analysis to inform strain engineering has been proposed^3,18–20^, and QTL mapping was previously used with marker assisted breeding for strain development^21^. The value for QTL analysis in strain engineering is sometimes evident in routine follow-up experiments using deletions, allele^†^ replacements, or allele complementation to confirm QTLs; the resulting strains occasionally display slightly improved phenotypes^18–24^. To our knowledge, however, no one has used QTL analysis to engineer a strain with a phenotype superior to both parents.

Engineering of complex traits can be hindered by unanticipated epistasis among otherwise beneficial alleles^19,23,25^. Epistasis in which the phenotypic effect of one or both alleles is inverted by genetic interaction^26,27^, when unanticipated, is problematic for strain development (e.g. two alleles, independently beneficial, produce an overall detrimental effect together). By contrast, combining independently beneficial alleles having unanticipated epistasis that only magnifies or mutes the expected effect^28^ is not problematic; the net effect of combining such alleles will be no less than the benefit independently gained from the larger effect allele. The existence of transgressive segregation suggests that engineering a superior strain should be possible by constructing the correct multi-locus genotype^2,29,30^; however, predicting this genotype is a major challenge and such strains are usually isolated by high-throughput screening and selection rather than by rational strain engineering^2–4^. Therefore, a general approach is needed to identify genetic regions relevant for strain engineering and to predict which alleles should be used to avoid the introduction of unwanted epistatic interactions.

Here, we devised an integrated approach for QTL-guided metabolic engineering, and used our method to engineer hydrolysate tolerance in *S. cerevisiae*. Our methodology integrates high-resolution QTL mapping (Figure 1A), bulk Reciprocal Hemizygosity Analysis (bRHA) (Figure 1B), and a Cas9-mediated method for efficient allele replacements (Figure 1C-E). QTL mapping and bRHA are first used to identify functional genetic elements and to reveal the most promising combination of alleles for those elements to use in engineering a complex trait. This data provides a genetic blueprint to guide strain engineering, which can be efficiently performed using a dual-guide Cas9 method to rapidly replace many genetic variants (single nucleotide polymorphisms (SNPs), multiple nucleotide polymorphisms (MNPs), and insertions or deletions (INDELs)) in large, highly homologous, genomic regions. Our approach does not require understanding the biological mechanism(s) underlying each genetic element, and should be generalizable to engineering any complex trait in *S. cerevisiae*, as well as other organisms.

**Figure 1.**
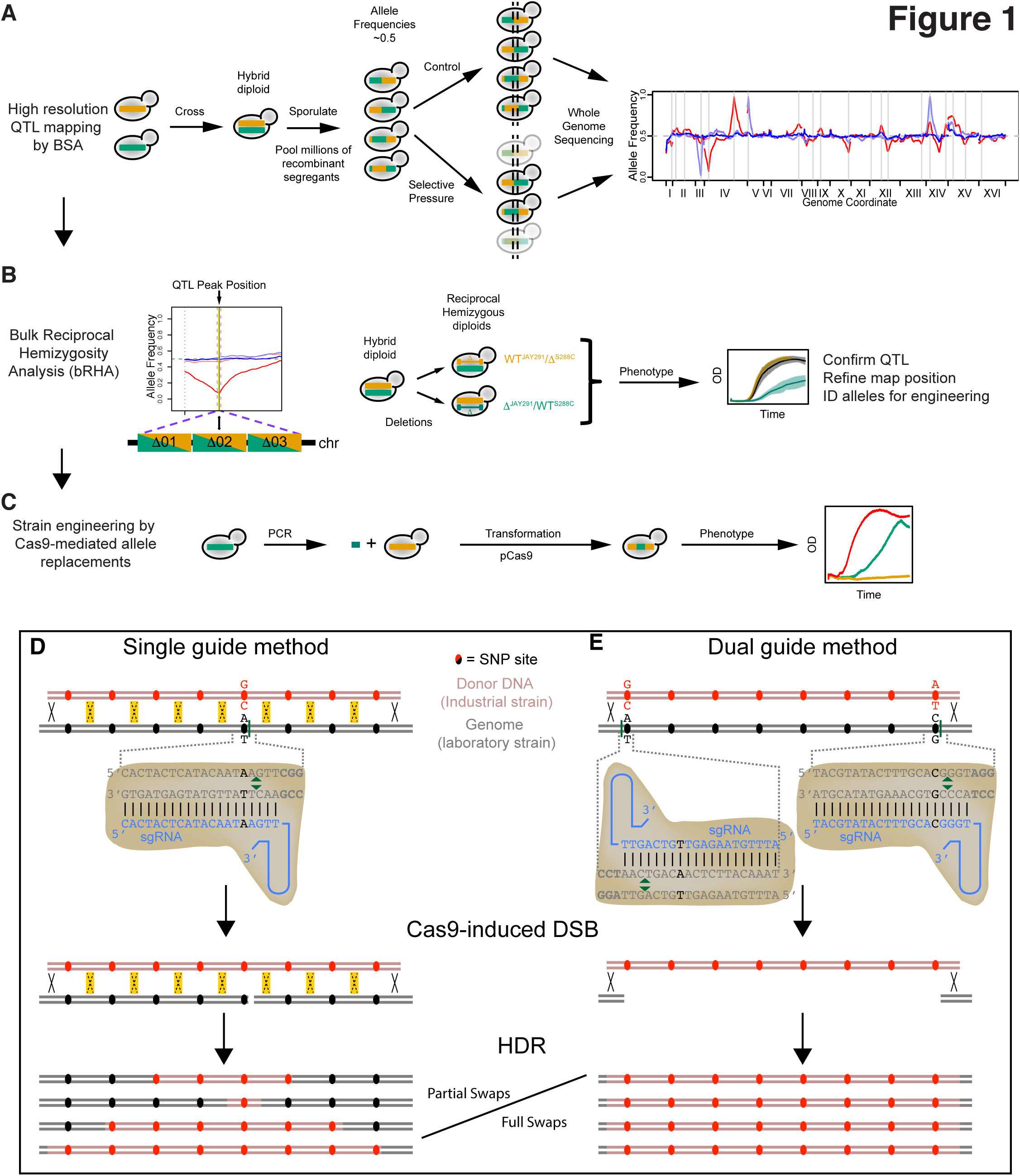
An integrated approach for QTL-mediated metabolic engineering of complex traits. (A) A bulk segregant analysis (BSA) approach for high-resolution QTL mapping is used to map QTLs (adapted from Parts *et al*^16^). The favorable QTL allele in the population is assessed from the vectored SNP allele frequency change at each QTL (see Fig 2A legend for details of the Allele Frequency vs Genome Coordinate plot). (**B**) Bulk Reciprocal Hemizygosity Analysis (bRHA) is used to confirm and refine the QTL map position for each QTL. Additionally, the phenotype of each reciprocal hemizygous strain is compared to the wild type (WT) hybrid diploid strain to determine whether the allele favorability manifests through a beneficial or detrimental mode of action. Allele favorability information from the QTL map and bRHA are used to create a genetic blueprint for strain engineering. (**C**) Strain construction using Cas9-mediated bulk variant replacements to combine bulk variant regions from different strains, according to the genetic blueprint. The desired bulk variant region is PCR amplified from the donor strain and co-transformed into the host strain with a plasmid expressing Cas9 (pCas9) and one or more sgRNAs. (**D-E**) Cas9 is directed to generate double strand breaks (DSBs) in the host chromosome within the intended bulk variant replacement region (green triangles indicate Cas9 cleavage sites). Variant sites (black text) within the Cas9-targeting sequences (protospacer + PAM) protect the donor DNA from Cas9 cleavage. The donor DNA is used in homology directed repair (HDR) of the DSBs. In the single-guide version diagrammed in **D**, only one Cas9-induced DSB is made. Here, resolution of the Holiday (i.e. recombination) junctions can occur throughout the highly homologous bulk variant replacement region (yellow-highlighted X symbols), resulting in various partial-region replacement events. In the dual-guide approach diagrammed in **E**, Cas9 is directed to make DSBs at two sites, promoting removal of the host bulk variant regions and recombination between the distal ends of the donor DNA.(A) A bulk segregant analysis (BSA) approach for high-resolution QTL mapping is used to map QTLs (adapted from Parts *et al*^16^). The favorable QTL allele in the population is assessed from the vectored SNP allele frequency change at each QTL (see Fig 2A legend for details of the Allele Frequency vs Genome Coordinate plot). (**B**) Bulk Reciprocal Hemizygosity Analysis (bRHA) is used to confirm and refine the QTL map position for each QTL. Additionally, the phenotype of each reciprocal hemizygous strain is compared to the wild type (WT) hybrid diploid strain to determine whether the allele favorability manifests through a beneficial or detrimental mode of action. Allele favorability information from the QTL map and bRHA are used to create a genetic blueprint for strain engineering. (**C**) Strain construction using Cas9-mediated bulk variant replacements to combine bulk variant regions from different strains, according to the genetic blueprint. The desired bulk variant region is PCR amplified from the donor strain and co-transformed into the host strain with a plasmid expressing Cas9 (pCas9) and one or more sgRNAs. (**D-E**) Cas9 is directed to generate double strand breaks (DSBs) in the host chromosome within the intended bulk variant replacement region (green triangles indicate Cas9 cleavage sites). Variant sites (black text) within the Cas9-targeting sequences (protospacer + PAM) protect the donor DNA from Cas9 cleavage. The donor DNA is used in homology directed repair (HDR) of the DSBs. In the single-guide version diagrammed in **D**, only one Cas9-induced DSB is made. Here, resolution of the Holiday (i.e. recombination) junctions can occur throughout the highly homologous bulk variant replacement region (yellow-highlighted X symbols), resulting in various partial-region replacement events. In the dual-guide approach diagrammed in **E**, Cas9 is directed to make DSBs at two sites, promoting removal of the host bulk variant regions and recombination between the distal ends of the donor DNA.

## Results & Discussion

### High-resolution QTL mapping of hydrolysate tolerance identifies seventeen QTLs

To identify genetic elements as potential parts for engineering hydrolysate tolerance in *S. cerevisiae*, we performed a Bulk Segregant Analysis (BSA) using a methodology we derived from three previous high-resolution QTL mapping studies^16,17,31^. We chose two haploid parent strains (JAY291 and S288C) for our QTL mapping cross that are genetically diverged^32^ (see also **Supporting Information**), and have distinct hydrolysate tolerance phenotypes (**Figure S1**). JAY291 is derived from an industrial strain, PE-2, that is widely used for sugarcane ethanol production in Brazil, and matches the industrial PE-2 strain in ethanol production^6,32,33^. S288C is a well-characterized laboratory strain that has a high-quality, well-annotated genome sequence^34^.

First, we generated a large pool of ~108 million haploid recombinant segregants^17^ (see also Methods). We then enriched for hydrolysate-tolerant segregants by growing this pool in an anaerobic continuous-culture fermentation system using defined media supplemented with 30% (vol/vol) dilute-acid treated *Miscanthus x giganteus* plant hydrolysate. We estimated population SNP allele frequencies, identified genomic regions with genetic variation responsible for the phenotypic variation, and determined the population-favorable QTL allele at those regions using a combination of published analysis methods^16,31,35^ (see also Methods). In total, we identified 17 QTLs (Figure 2A, Table 1 and **Table S1**), confirming that hydrolysate tolerance is a complex genetic trait^12,36,37^. We observed favorable QTL alleles of both parental ancestries (eight JAY291 and nine S288C alleles) (Figure 2A and Table 1). The resolution of the identified QTLs, defined as the 95% confidence interval (CI) of the predicted QTL location, ranged from ~6 kb to 732 kb; therefore, further QTL map-refinement was necessary to enable efficient strain engineering.

**Figure 2.**
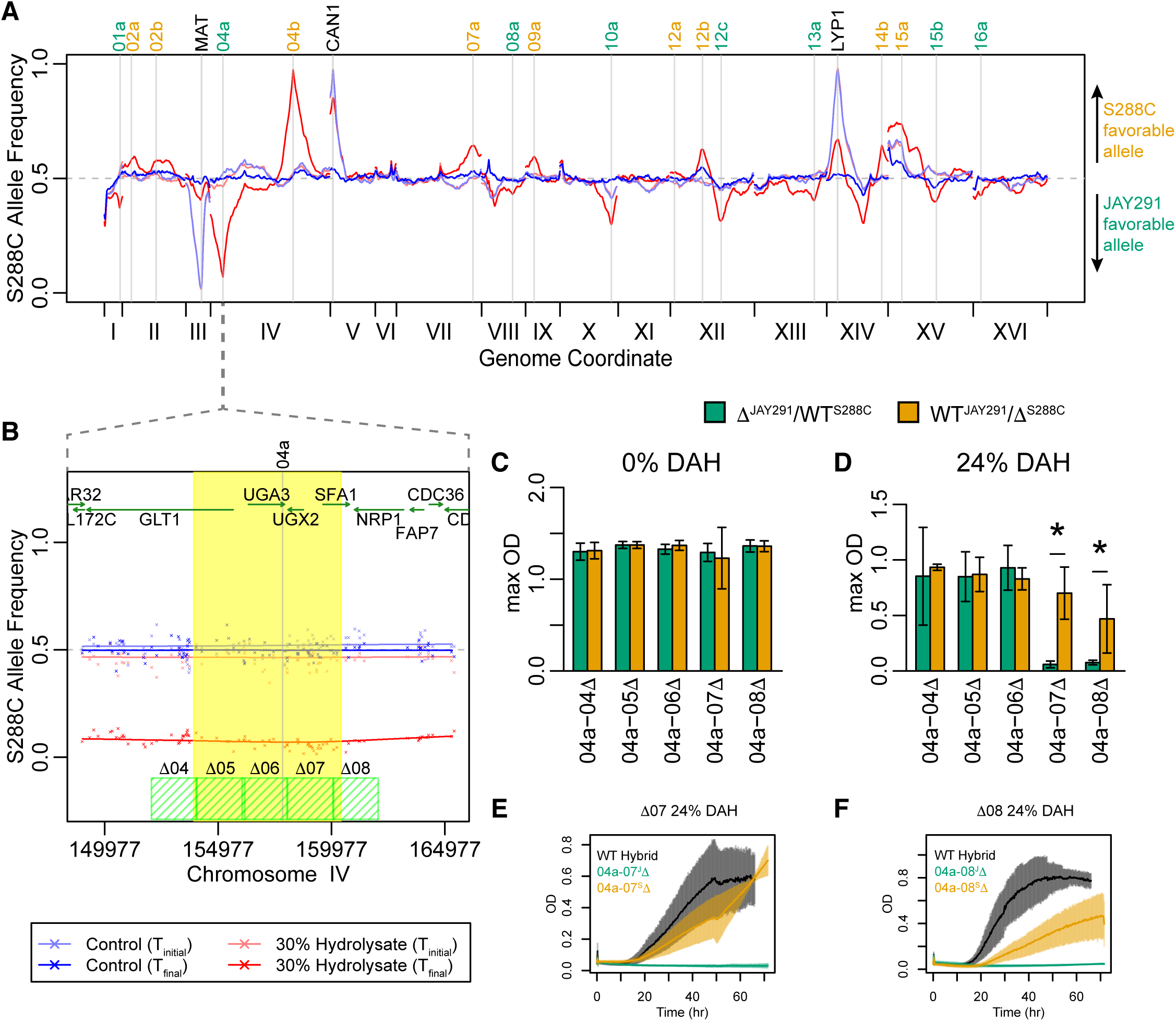
Creating a “genetic blueprint” from the QTL map and bRHA. (**A**) High-resolution mapping of 17 hydrolysate-tolerance QTLs. The inferred S288C SNP allele frequencies for the populations cultured in 0% and 30% (vol/vol) *Miscanthus x giganteus* dilute acid hydrolysate (DAH) at the time of inoculation (T_initial_, light blue and light red lines, respectively) and after 75 population doublings (T_final_, dark blue and dark red lines, respectively) are plotted against genome position for all chromosomes. Positions of the best predictions for the 17 QTLs, and the *MAT*, *CAN1*, and *LYP1* loci which served in segregant pool generation (see Methods), are shown (vertical gray lines). (**B**) An ~15 kb window surrounding QTL 04a, enlarged from panel **A**. The inferred and observed SNP allele frequencies (lines and “x” symbols, respectively, color-coded as in **A**), 95% confidence interval for predicted position of QTL 04a (yellow shaded region), gene ORFs (green arrows), and bRHA deletion regions (green hashed boxes) are shown. (**C**-**D**) QTL 04a map position is confirmed and refined by bRHA to the Δ07-Δ08 region. Reciprocal hemizygous strains were constructed by deleting either the JAY291 allele or S288C allele for each region corresponding to the green hashed boxes in **B**. Mean maximum culture density (max OD) is plotted for reciprocal hemizygous strains deleted for the JAY291 allele (Δ^JAY291^/WT^S288C^, green bars) and the S288C allele (WT^JAY291^/Δ^S288C^, orange bars) grown anaerobically in microplates in synthetic media lacking hydrolysate (**C**), or synthetic media supplemented with 24% (vol/vol) DAH (**D**). Error bars represent 95% confidence intervals calculated from the t-distribution (n ≥ 3 biological replicates for each strain in each condition). Reciprocal hemizygous strains with statistically significant fitness differences within the strain pair are indicated (*; p<0.05 by Welch’s t-test). (**E**-**F**) A favorable JAY291 allele at QTL 04a is beneficial to the wild type (WT) hybrid diploid strain. The growth curves for the WT hybrid diploid strain and reciprocal hemizygous strains for the Δ07 (**E**) and Δ08 (**F**) regions, grown in synthetic media supplemented with 24% (vol/vol) DAH. Traces of mean culture densities (OD) are plotted as a function of time. Error bars indicate one standard deviation. Means and standard deviations reported for the WT hybrid diploid were calculated using only trials performed in the same microplates as the reciprocal hemizygous strains appearing in the plots (n ≥ 3 biological replicates for each strain in each condition).

**Table 1.** Allele favorability and effect for hydrolysate tolerance QTLs

| QTL Name | Population Favorable Allele | bRHA Favorable Allele | JAY291 allele effect in hybrid diploid | S288C allele effect in hybrid diploid | Region(s) Tested by bRHA | Region(s) Confirmed by bRHA |
| --- | --- | --- | --- | --- | --- | --- |
| 01a | JAY291 | S288C | Neutral | Beneficial | 2 | 2 |
| 02a | S288C | not determined | not determined | not determined | 1,2,3 | none |
| 02b | S288C | not determined | not determined | not determined | 2 | none |
| **04a** | **JAY291** | **JAY291** | **Beneficial** | **Beneficial** | **2kb: {4,5,6,7,8,9}** | **2kb: {7,8}** |
| **04b** | **S288C** | **S288C** | **Detrimental** | **Beneficial<sup>†</sup>** | **2kb: {1,2,3,4,5}** | **2kb: {2,3}** |
| **07a** | **S288C** | **S288C** | **Neutral** | **Beneficial** | **1,2,3** | **1** |
| 08a | JAY291 | not tested | not tested | not tested | none tested | none tested |
| 09a | S288C | not determined | not determined | not determined | 2 | none |
| **10a** | **JAY291** | **JAY291** | **Beneficial** | **Neutral** | **1,2,3** | **1,2** |
| 12a | S288C | JAY291 | Beneficial | Detrimental | 1,2,3 | 3 |
| 12b | S288C | JAY291 | Neutral | Detrimental | 1,2,3 | 1 |
| **12c** | **JAY291** | **JAY291** | **Beneficial** | **Neutral** | **1** | **1** |
| 13a | JAY291 | not determined | Beneficial | Beneficial | 1,2,3 | 1 |
| **14b** | **S288C** | **S288C** | **Detrimental** | **Neutral** | **1,2,3** | **1** |
| 15a | S288C | not determined | Beneficial | Beneficial | 2 | 2 |
| 15b | JAY291 | JAY291 | Neutral | Detrimental | 2 | 2 |
| 16a | JAY291 | S288C | Neutral | Beneficial | 2 | 2 |

<sup>†</sup>Also beneficial for general fitness

### Creating a “genetic blueprint” for strain engineering using the QTL map and bRHA

Because QTL 04a was identified with high resolution (95% CI of 6.5 kb), we scanned across the entire 95% CI using bRHA (Figure 2B, subregions Δ04 thru Δ08). We constructed six reciprocal hemizygous strain pairs and phenotyped them for hydrolysate tolerance using a microplate growth assay. No fitness difference was observed within reciprocal pairs in the no-hydrolysate control condition (Figure 2C and **Figure S12B**). By contrast, deleting the JAY291 allele of either subregion 04a-Δ07 or 04a-Δ08 more strongly decreased fitness in 24% (vol/vol) hydrolysate than deleting the S288C allele (Figure 2C-F). The remaining reciprocal pairs experienced little to no relative fitness difference in hydrolysate (Figure 2D and **Figure S12C** and **D**, panels Δ04, Δ05, and Δ06. Thus, using bRHA, we confirmed the presence of a favorable JAY291 allele underlying QTL 04a and refined its position to a 4 kb genomic region. The fact that deleting either allele of 04a-Δ07 and 04a-Δ08 reduced hydrolysate tolerance relative to the wild type (WT) hybrid diploid (Figure 2E and F), reveals haploinsufficiency for QTL 04a. This result indicates both alleles of 04a are beneficial to the hybrid diploid strain, albeit the JAY291 allele to a larger degree.

We systematically used bRHA to analyze a 5-15 kb region centered around 15 additional predicted QTLs (**Figures S9-25**, see also **Supporting Information**). In nearly all cases (13 of 16), we confirmed the presence of hydrolysate tolerance alleles and refined their map position.

This is not to say we identified all phenotypically relevant genetic variants underlying a particular QTL, as other causal variants may lie in the untested regions of the predicted QTLs. We also classified each allele’s effect (beneficial, detrimental, or neutral) and favorability (JAY291 or S288C) pertaining to hydrolysate tolerance in the hybrid diploid (Table 1, see also **Supporting Information**). We identified four JAY291 QTL alleles (04a^J^, 10a^J^, 12c^J^ and 15b^J^) that were favorable in both the segregant population and the hybrid diploid. Three of these JAY291 QTL alleles (04a^J^, 10a^J^, and 12c^J^) were beneficial to the hybrid diploid strain, whereas for QTL 15b, the JAY291 QTL allele was neutral; the favorability at 15b stemmed from a detrimental effect of the S288C QTL allele, which was only very small. There were also three S288C QTL alleles (04b^S^, 07a^S^, and 14b^S^) that were favorable in both the segregant population and the hybrid diploid. In four cases (QTLs 01a, 12a, 12b, and 16a), QTL allele favorability assessments by BSA and bRHA were discordant. These discrepancies could result from two different, tightly linked QTLs with opposite favorable alleles underlying a single predicted QTL region^24,25,38^ or strain-dependent genetic interactions that cause an allele effect inversion (epistasis)^26–28,39–41^. Either tightly linked QTLs or allele effect inversion could confound strain design and engineering; therefore, we focused on QTLs 04a, 04b, 07a, 10a, 12c, and 14b (**Supporting Information**) that were unlikely to be problematic.

### Efficient allele replacements in large regions of highly homologous DNA using Cas9

To facilitate engineering of complex phenotypes, we implemented two Cas9-mediated strategies (single- and dual-guide) for efficient bulk variant replacements (Figure 1D and E and **Supporting Information**). Simultaneous replacement of many variants in large regions of highly homologous DNA (such as between JAY291 and S288C) is challenging and this step was a significant bottleneck in strain engineering. Although complete bulk variant replacements occurred, the single-guide method often resulted in a partial outcome, where only a subset of the targeted variant sites was replaced (Figure 1D and **Figure S3A** and **B**). However, we took advantage of these instances to perform ultra-fine QTL mapping, which provided biological insight into a mechanism of hydrolysate tolerance (see below). By contrast, the dual-guide method excelled at producing full bulk variant replacements and was routinely used to substitute between 9 and 40 variants across ~5 kb regions during strain engineering (Figure 1E, **Figure S3C**, and unpublished observations).

### Defining a functional part for engineering hydrolysate tolerance: 04a-07

To determine whether the bRHA-confirmed region 04a-07 (Figure 2 and **Figure S12**), could serve as a functional part for engineering hydrolysate tolerance, we performed a bulk variant replacement for 04a-07. We transferred the JAY291 variants for this region (28 SNPs and one 1 bp INDEL) into S288C using our single-guide Cas9 method (Figure 3). These variant substitutions in this S288C+04a-07^J^ strain had no effect on general fitness (**Figure S4B**), but were sufficient to increase hydrolysate tolerance in S288C (Figure 3B and **C** and **Figure S4G** and **L**). Thus, this S288C to JAY291 bulk variant substitution (04a-07^S→J^) is a functional genetic part replacement that improves hydrolysate tolerance in S288C. The process of combining information from BSA and bRHA, and performing Cas9-mediated bulk variant replacements, demonstrated here for 04a-07, exemplifies our integrated method for QTL-guided metabolic engineering.

**Figure 3.**
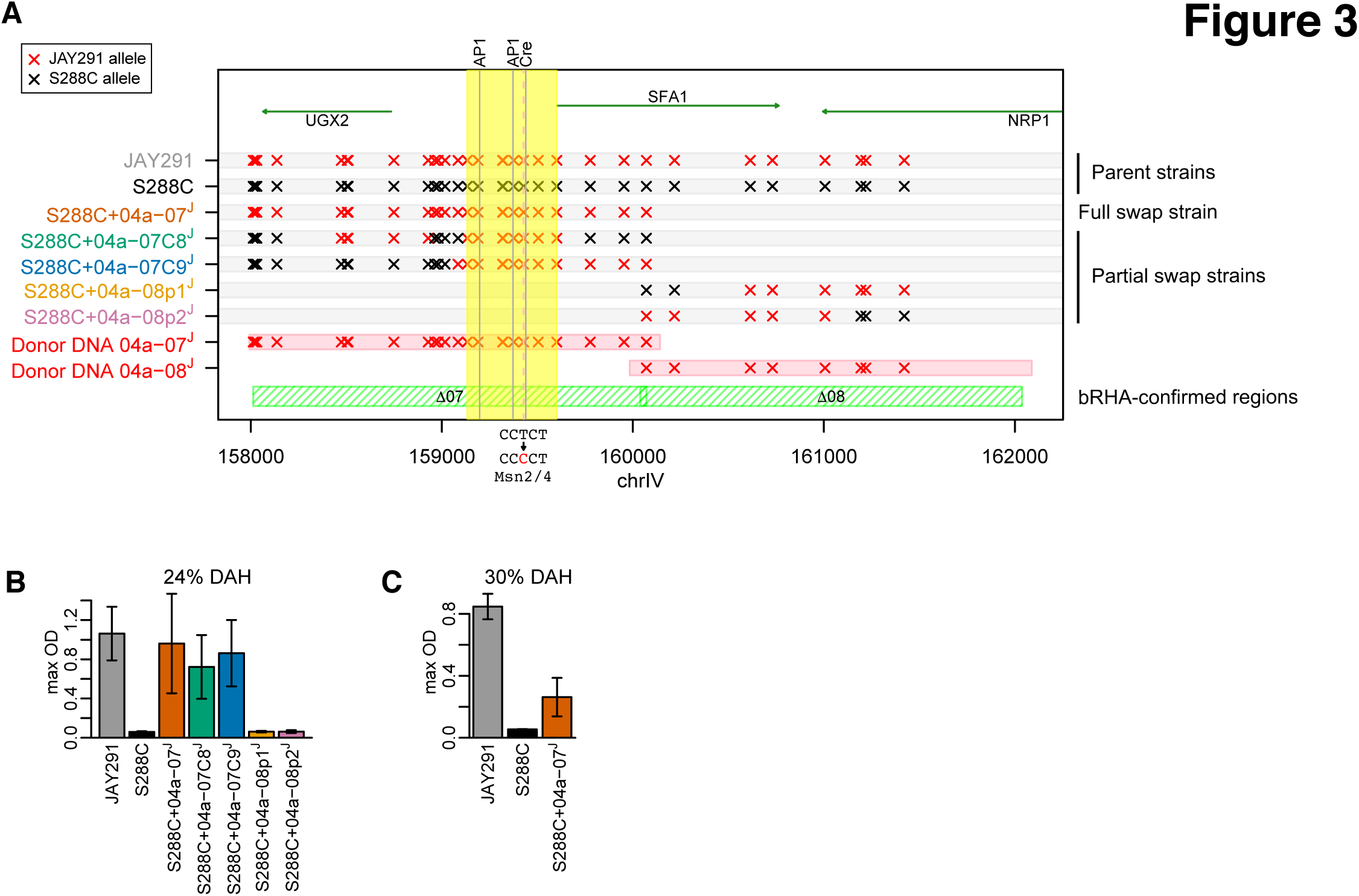
Analysis of Cas9-mediated partial allele replacement strains reveals variants in the *SFA1* promoter region important for hydrolysate tolerance. (**A**) Genotypes of five different full and partial allele replacement strains for QTL 04a, generated by the single-guide Cas9-mediated allele replacement method. Red and black “x” symbols signify JAY291 and S288C alleles, respectively, at known SNP sites. Donor DNA (red bars, “Donor DNA 04a-07^J^” and “Donor DNA 04a-08^J^”), containing the JAY291 alleles at variant sites within bRHA-confirmed regions (green hashed bars, Δ07 and Δ08), were generated by PCR and used to replace S288C alleles in the S288C genome. Transformants were Sanger sequenced to determine the genotype at each variant site. Genotypes at SNP sites for JAY291 and S288C are provided as a reference. Gene ORFs (green arrows) and known *SFA1*-promoter elements (AP-1 binding sites at −409/−403 and −234/−228, Cre element at −168/−161^45^, vertical gray lines), and a T→C SNP at −176 that creates an Msn2/4 binding site^46^ (vertical dashed red line) are also indicated. The region spanning substituted variant sites common in the three improved strains is highlighted yellow. (**B-C**) Replacing S288C variants with JAY291 variants in region 04a−07 improves hydrolysate tolerance. Mean maximum culture densities (max OD) are plotted for the indicated strains grown anaerobically in microplates in synthetic media supplemented with either 24% (**B**), or 30% (vol/vol) dilute acid hydrolysate (DAH) (**C**). Error bars represent 95% confidence intervals calculated from the t-distribution. Means and confidence intervals reported for JAY291 and S288C were calculated using only trials performed in the same microplates as the allele replacement strains appearing in the same plots (n ≥ 3 biological replicates for each strain in each condition).

### Cas9-mediated partial allele replacements facilitate ultra-fine mapping of QTL parts

We used the partial substitution strains for 04a−07 and 04a−08, generated by our single-guide Cas9-mediated bulk variant replacement method (Figure 3A and **Figure S3A** and **B**) to elucidate the biological mechanism underlying QTL 04a. Our bRHA results for 04a-Δ07 and 04a-Δ08, that each partially delete *SFA1* (Figure 2B and **D-F**), implicate this gene. *SFA1* encodes a dehydrogenase that catabolizes growth inhibitors in hydrolysate^42–44^. The observed haploinsufficiency (Figure 2E-F) could stem from decreased Sfa1p activity, but whether the favorable JAY291 variants result in more protein, or higher intrinsic enzymatic activity, is indeterminable by bRHA alone.

The partially substituted 04a−07^J^ strains were more tolerant to hydrolysate than S288C (Figure 3B and **Figure S4H** and **I**), whereas the partial 04a−08^J^ replacement strains were not (Figure 3B and **Figure S4J** and **K**). Comparing the common variant substitutions in the improved strains reveals that variants critical for improved hydrolysate tolerance reside within the *UGX2–SFA1* intergenic region (Figure 3). Intergenic SNPs at positions −414, −227, and −176, relative to *SFA1* +1, are nearby known *SFA1* promoter elements^45^ (Figure 3A). Furthermore, the T to C SNP at −176 creates an Msn2/4p binding site that is known to increase *SFA1* expression^46^. The hydrolysate-tolerance variant positions suggest differential *SFA1* expression, rather than intrinsic enzyme activity, contribute to the phenotypic difference between S288C and JAY291.

### A 5 kb region at QTL 10a is identified as a part for hydrolysate tolerance engineering

Our bRHA of two 5 kb regions (10a-Δ01 and 10a-Δ02) confirmed beneficial JAY291 variants in QTL 10a (**Figure S17** and **Supporting Information**). Therefore, we performed bulk variant replacements in these regions using our dual-guide Cas9 method, generating two completely substituted strains (S288C+10a−01^J^ and S288C+10a−02^J^) (**Figure S5A**). Only the 10a−02^S→J^ replacement had a small, but statistically significant, effect on hydrolysate tolerance (Figure 4A and **Figure S5B** and **C**). Thus, without any knowledge of the mechanism, or having identified the causative gene, we defined a second functional bulk variant replacement (10a−02^S→J^) for strain engineering.

**Figure 4.**
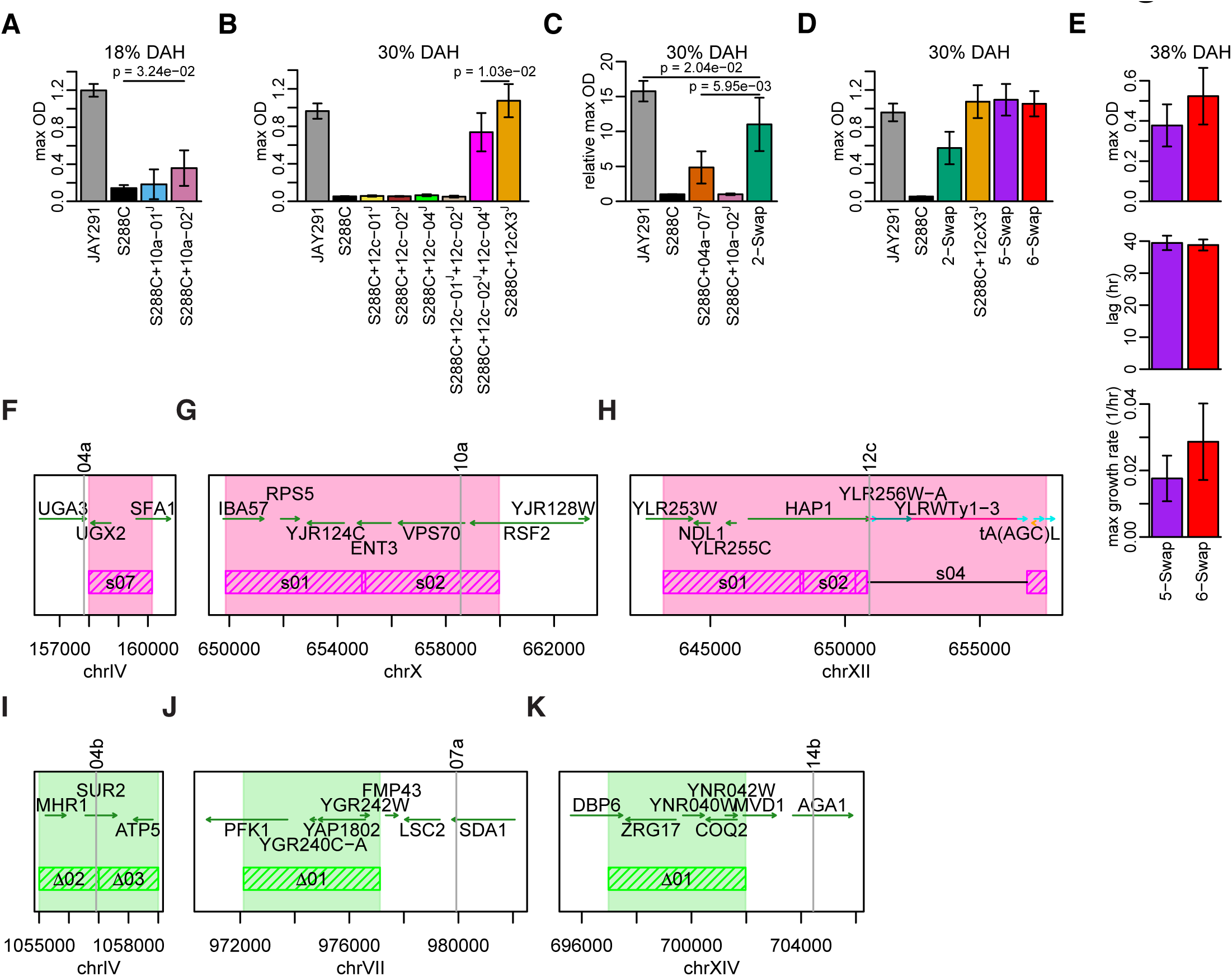
Combining allele replacements provides incremental improvements in hydrolysate tolerance. (**A**) The 10a−02^J^ allele replacement improves hydrolysate tolerance of S288C. Mean maximum culture densities (max OD) are plotted for the indicated strains grown anaerobically in microplates in synthetic media supplemented with 18% (vol/vol) dilute acid hydrolysate (DAH).(B) Substituting multiple variants in *HAP1* at QTL 12c is necessary to improve hydrolysate tolerance in S288C. Mean maximum culture densities (max OD) are plotted for the indicated strains grown anaerobically in microplates in synthetic media supplemented with 30% (vol/vol) DAH. (**C**) Positive epistasis exists between the 04a−07^J^ and 10a−02^J^ alleles: the relative fold change between S288C and the 2-swap strain (10.01) is greater than the sum of the fold changes for S288C+04a−07^J^ (3.85) and S288C+10a−02^J^ (0.01), relative to S288C. Relative maximum culture densities (relative max OD) were calculated for the indicated strains grown anaerobically in microplates with 30% (vol/vol) DAH. (**D**) The 5-swap and 6-swap strains are more tolerant to hydrolysate than the 2-swap strain. Mean maximum culture densities (max OD) for the indicated strains grown anaerobically in microplates in synthetic media supplemented with 30% (vol/vol) hydrolysate. (**E**) Maximum culture density (max OD), lag phase duration (lag), and max growth rate estimates from fitted logistic growth curve models for the 5-swap and 6-swap strains at 38% (vol/vol) DAH. In **A-E**, error bars represent 95% confidence intervals calculated from the t-distribution. Black horizontal bridging lines represent statistically significant differences; p-values were calculated using Welch’s t-test. The means, confidence intervals, and p-values were calculated from three or more biological replicates for each strain and condition. Values reported for JAY291 and S288C were calculated using only trials performed in the same microplates as the allele replacement strains appearing in the same plot. (**F-K**) Genomic regions for QTLs with favorable JAY291 and S288C alleles, predicted to be compatible for engineering hydrolysate tolerance. Substitution regions (**F-H**, pink hashed bars and highlighted regions) are shown for QTLs with favorable JAY291 alleles, and bRHA-confirmed regions (**I-K**, green hashed bars and highlighted regions) are shown for QTLs with favorable S288C alleles. Best predicted position for each QTL is (vertical gray lines), protein-coding gene ORFs (green arrows), tRNA genes (orange arrows), retrotransposons (pink), long terminal repeats (cyan), and transposable element genes (dark cyan) are shown.

### Allele replacements at QTL 12c indicate *HAP1* is a part for hydrolysate tolerance engineering

Our bRHA of the 5 kb 12c-Δ01 region, just upstream of the predicted QTL 12c peak, confirmed beneficial JAY291 variants for hydrolysate tolerance. However, a Ty1 element in S288C that is absent in JAY291^32^, prevented us from scanning across this peak (**Figure S20** and **Supporting Information**). In S288C, this Ty1 element alters the C-terminus of the pleiotropic Hap1p transcription factor^32,47^. We therefore designed bulk variant replacements for three QTL 12c subregions (12c−01, 12c−02 and 12c−04, **Figure S6A**). The 12c−02^S→J^ and 12c−04^S→J^ replacements queried all remaining *HAP1* variants not covered by 12c−01^S→J^, including the Ty1 element and 11 additional variants in 676 bp of 3’ UTR (relative to *HAP1* in JAY291)^48^. None of the single region replacements, or the combination of 12c−01^S→J^ and 12c−02^S→J^ together, had any effect on hydrolysate tolerance (Figure 4B and **Figure S6C-F** and **J**). By contrast, combining 12c−02^S→J^ and 12c−04^S→J^ greatly improved hydrolysate tolerance (Figure 4B and **Figure S6G-J**), and replacing all three regions together (12c−01+12c−02+12c−04, herein referred to as 12cX3) provided additional improvement (Figure 4B and **Figure S6H-J**, S288C+12cX3^J^). These results indicate multiple *HAP1* variants are important for hydrolysate tolerance, and reveal positive epistasis between *HAP1* variants. Furthermore, these results identify the 12cX3^S→J^ substitution as a third part replacement for improving hydrolysate tolerance in S288C.

### QTL-guided metabolic engineering results in a superior phenotype

Because both parents could provide favorable alleles (Figure 2, **Figures S12-14**,**17**,**20**,**22**, and Table 1), we reasoned that it could be possible to engineer S288C with greater hydrolysate tolerance than JAY291 by replacing, in the correct combination, multiple S288C bulk variant regions with JAY291 versions. Thus far, we identified functional bulk variant replacements, substituting JAY291 variants into S288C at QTLs 04a, 10a, and 12c (Figure 4F-H), that individually improved hydrolysate tolerance (Figures 3 and 4A and B). Based on our BSA and bRHA data, we predicted that these substitutions would confer greater hydrolysate tolerance together than individually (see **Supporting Information**). To test this idea, we began combining the 04a−07^S→J^, 10a−02^S→J^, and 12cX3^S→J^ bulk variant replacements in S288C and phenotyped the resulting strains for hydrolysate tolerance. We screened these strains with our microplate growth assay to identify candidate strains for further testing in a more industrially relevant, but lower throughput, bioreactor format.

When we combined the 04a−07^S→J^ and 10a−02^S→J^ replacements in S288C, the resulting strain (called 2-swap) was more tolerant to hydrolysate than S288C with either single bulk variant replacement. Moreover, we observed positive epistasis between 04a−07^J^ and 10a−02^J^ (Figure 4C and **Figure S7A**). We then combined the 12cX3^S→J^, 04a-07^S→J^ and 10a−02^S→J^ replacements into a single strain (called 5-swap), and compared its hydrolysate tolerance to the 2-swap and S288C+12cX3^J^ strains. We generated an additional strain (called 6-swap) by also including the 10a−01^S→J^ replacement; although the 10a−01^S→J^ replacement did not improve hydrolysate tolerance on its own (Figure 4A), our bRHA results for 10a-Δ01 (**Figure S17** and **Supporting Information**), and the positive epistasis within QTL 12c (Figure 4B and **Figure S7B**) motivated us to test this 6-swap genotype. At 30% (vol/vol) hydrolysate, the 5-swap, 6-swap, and S288C+12cX3^J^ strains were more tolerant than the 2-swap strain, but indistinguishable from each other (Figure 4D). At 38% (vol/vol) hydrolysate, however, we observed higher mean maximums for culture density and growth rate for the 6-swap strain than the 5-swap strain (Figure 4E and **Figure S7C**) at low statistical significance (p=0.078 and p=0.109, respectively). Because error in our microplate format increases with hydrolysate concentration (see **Figure S8A-D** for example), we transitioned to the bioreactor format for further testing at higher hydrolysate concentrations.

We tested the 6-swap strain against S288C, S288C+12cX3^J^, and JAY291, in anaerobic batch fermentations containing 40% (vol/vol) hydrolysate in bioreactors (Figure 5). Because ethanol production is tightly coupled with anaerobic growth in *S. cerevisiae*^49^, improving hydrolysate tolerance could also improve ethanol production. We therefore also assessed ethanol production and sugar utilization. Although these strains were equally fit in our microplate growth assays lacking hydrolysate, bioreactor growth-profile comparisons show that the 6-swap strain is superior to the S288C, S288C+12cX3^J^, and JAY291 strains in 40% (vol/vol) hydrolysate (Figure 5 and **Figure S8**). While maximum culture densities, total glucose consumption, and final ethanol titer were roughly equivalent between JAY291 and the 6-swap strain, our engineered strain had a shorter lag phase, higher maximal growth rate, and began producing ethanol earlier (Figure 5 and **Figure S8E-H**).

**Figure 5.**
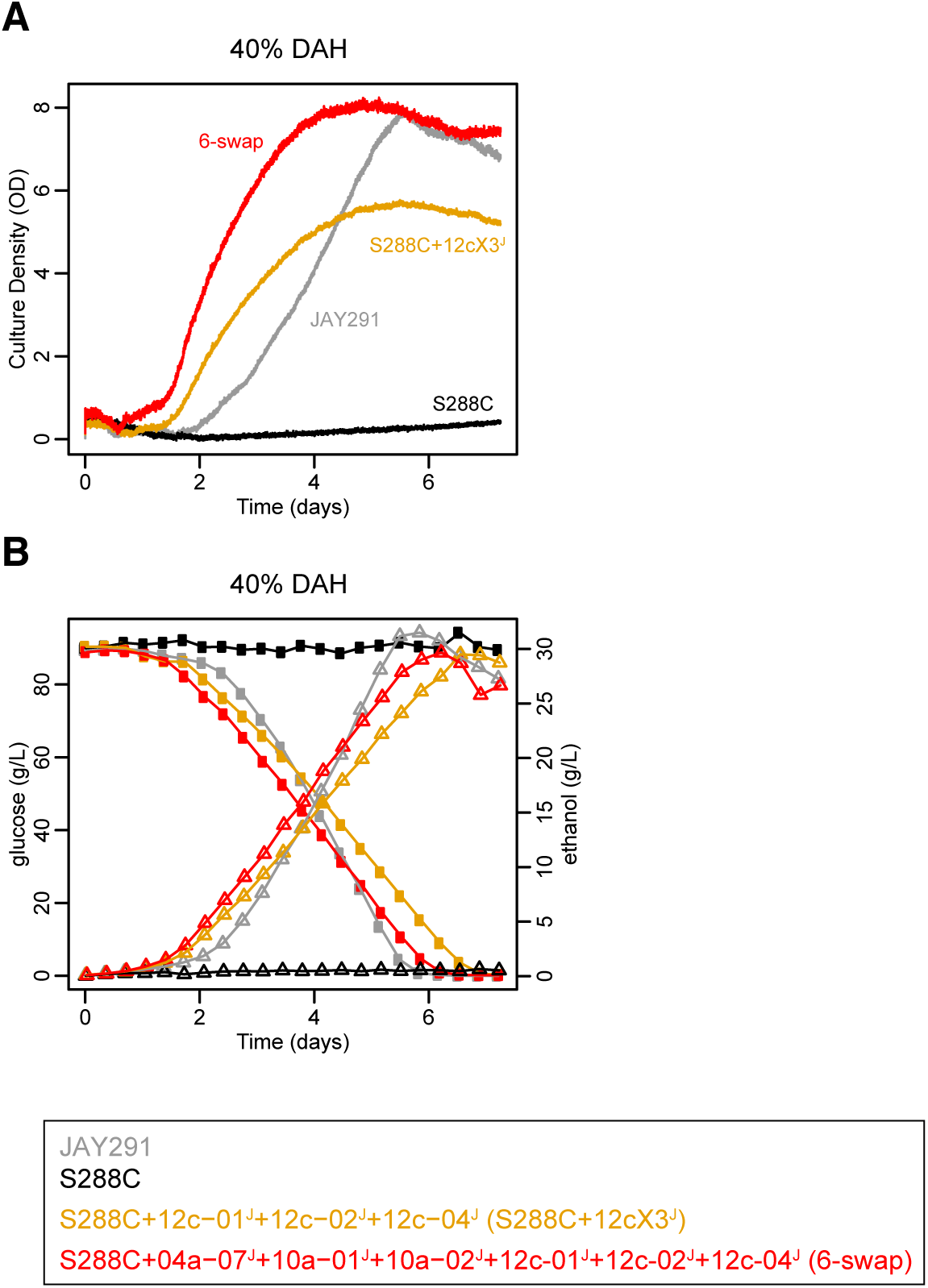
QTL-guided metabolic engineering results in a strain with superior hydrolysate tolerance phenotype. (**A-B**) Anaerobic batch fermentation profiles using a DASGIP bioreactor in synthetic media supplement with 40% (vol/vol) dilute acid hydrolysate (DAH) for the S288C laboratory strain, JAY291 industrial strain, and two engineered strains, “S288C+12cX3^J^” and “6-swap”. The genotype of each strain is shown in the legend. Culture density (OD) (**A**, unmarked solid lines), glucose (g/L) (**B**, circles), and ethanol (g/L) (**B**, triangles) values are plotted as a function of fermentation time (days) for one representative experiment. Additional replicate experiments are shown in **Figure S8**.

## Conclusions

Engineering complex, polygenic traits for industrial applications is challenging. Here we present the first demonstration, to our knowledge, of successfully implementing QTL analysis for the design and construction of a strain with a phenotype superior to either parent (Figure 5A and **Figure S8E-H**, top panels). Our integrated approach reveals functional genetic elements that can serve as parts for building a complex trait. By replacing variants in S288C at QTLs 04a, 10a, and 12c (Figure 4F-H) with the favorable variants from JAY291, while leaving the favorable S288C variants at QTLs 04b, 07a, and 14b unchanged (Figure 4G-K), we were able to improve hydrolysate tolerance in S288C to a level beyond JAY291 (Figure 5A and **Figure S8E-H**, top panels).

The success of QTL-guided complex trait engineering is dependent on creating a high-resolution QTL map, not the trait being mapped, nor the mapping method. An alternative to BSA is Independent Segregant Analysis (ISA), in which each recombinant segregant is phenotyped and genotyped individually rather than phenotyping and genotyping at the population level. Though ISA is generally more laborious and expensive than BSA, this does not preclude its use; ISA can be and has been used to map QTLs at high resolution^50,51^, including QTLs for non-selectable traits such as colony morphology ^50^. So given that high resolution QTL mapping is not dependent on phenotyping and genotyping at the population level, and has been successful for both fitness-based and non-fitness based traits, including non-selectable traits^16,17,50,51^, it is reasonable to expect that our approach can be generally applied to any complex trait.

A critical bottleneck in strain engineering is the ability to efficiently perform variant replacements between two *S. cerevisiae* strains. To overcome this problem, we implemented a dual-guide Cas9-mediated approach that was more efficient than standard technologies, and allowed us to engineer a strain where we substituted 109 variants across ~26.5 kb of DNA at three distinct genomic loci. The partial bulk variant replacements we observed using our single-guide method were not unexpected; resolution of the Holliday (i.e. recombination) junctions can occur throughout the highly homologous allele replacement region^52,53^, resulting in various partial-region replacement events (Figure 1D, yellow-highlighted X symbols, see also **Supporting Information**). By contrast, the dual-guide, Cas9-mediated strategy enables direct, simultaneous replacement of many genetic variants across large, highly homologous regions of DNA, and is currently the method of choice for our strain engineering projects.

Our approach does not require understanding the biology underlying the genetic parts, however, it can aid in elucidating the biological mechanisms, and facilitate alternative engineering approaches. In the course of our engineering, we identified *SFA1* and *HAP1* as causative genes in two QTLs and determined that important variants exist in the *SFA1* promoter region (Figure 4). Hap1p indirectly regulates *SFA1* via Hog1p^45^. Hap1p also binds the promoter of *COQ2*, located in the 14b-Δ01 subregion of QTL 14b (**Figure S22A**), and downregulates *COQ2* expression under hypoxic conditions^48^. It could be that *COQ2* variants are responsible for QTL 14b. Combining our BSA, bRHA, and variant detection data with known genetic and physical interaction networks suggests additional connections between candidate genes for future studies (unpublished observations). Exploring these connections, and additional QTL dissection with the single-guide Cas9 strategy, could help elucidate the biology underlying the trait, and reveal specific DNA and/or amino acid residues to target with additional engineering approaches.

Genetic interactions were present among the hydrolysate-tolerance QTLs. The causal variants in QTLs 04a, 10a, and 12c were not simply additive; we observed positive epistasis between QTLs 04a and 10a (Figure 4C and **Figure S7A**), and variants within QTL 12c displayed intragenic epistasis (Figure 4B and **Figure S6**). For at least four other QTLs (QTLs 01a, 12a, 12b, and 16a), the favorable QTL allele in the segregant pool was the opposite allele of that which was favorable in the hybrid diploid strain, suggesting alleles for these QTLs likely experience genetic interactions with one or more other loci (see **Supporting Information**). We used bRHA in conjunction with BSA to guide our strain engineering, translating QTL allele behaviors in a hybrid diploid strain and segregant population into predictions for their expected behavior in an engineered haploid S288C strain. The incremental improvements in hydrolysate tolerance achieved by stacking bulk variant replacements (Figures 3,4,5A) indicate there was no epistatic inversion of allele effect, as we predicted.

## Future Considerations

In this study, we used the first filial (F1) generation of a cross between two parents for QTL mapping; however, several variations of this BSA-based method have recently been reported. The use of an F12 advanced intercross population (AIP) was shown to improve map resolution, but required significantly more time and effort to prepare the mapping population^16^. High-resolution BSA-based QTL mapping studies have also utilized multiple parent strains, which increased the genetic and phenotypic diversity that could be assessed in a single study^54–56^. In particular, Cubillos and colleagues constructed a multi-parent AIP^55^, which produced more distinct genotypes than pair-wise crosses, even when multiple parents are used. Consequently, there is greater potential to detect loci whose effects are context-dependent (epistatic), and the increased genetic and phenotypic diversity in a single population make that population more applicable to multiple traits. Based on these studies, we envision that the use of a multi-parent AIP will be a valuable improvement to our existing approach.

Some QTL mapping methods, such as ISA, can reveal ideal multi-locus genotypes, but as mentioned above, are generally more labor-intensive and costly than BSA^15,50^. Thus, developing the ability to infer ideal multi-locus genotypes from BSA data will also enhance QTL-guided engineering. Determining the ideal genotype by trial and error quickly becomes unreasonable; the number of possible haploid genotypes, given two alleles per QTL, grows with the number of QTLs as a power of two. Previous work has suggested that it may be possible to infer epistasis from allele frequency flux in BSA data^16,35^. A complete picture of epistasis could drastically reduce the number of design-test-build cycles during strain engineering.

Systems and inverse metabolic engineering of polygenic, complex traits, which describe our QTL-based approach, have generally succeeded more than breeding and rational engineering^2^. Breeding in particular, typically produces intermediate phenotypes and improvements in one phenotype often come at the expense of another due to linkage and breeding’s reliance on sexual recombination^2,21,57^. Since breeding alone does not identify the important genetic determinants, the bred strain is a black box, and the phenotype cannot be reconstituted later if lost, or engineered into another strain. This same black box issue is true for genome shuffling, and others have suggested following genome shuffling with QTL analysis to overcome this issue^58^. Like breeding, our approach involves a genetic cross and exploits existing genetic diversity. Given that mating is one method used in genome shuffling, the work we describe here could also be considered genome shuffling followed by QTL analysis. By generating a QTL map to identify the genetic loci pertinent to engineering and create a blueprint for strain construction, and subsequently using targeted allele replacements to reconstruct an improved strain from identified parts, one may be able to overcome potential pitfalls faced during breeding and genome shuffling. With further development, QTL-guided host development may become a preferred approach.

The system we developed for QTL-guided complex trait engineering can be applied to both haploids and diploid organisms, and is not limited to selectable traits^16,17,50,51^. In addition, our approach can be integrated with other strain improvement methods, allowing it to be assimilated into a larger pipeline^1^. We expect our integrated approach for QTL-guided metabolic engineering to have a broad impact on host development for biomanufacturing a wide variety of products, such as therapeutics, renewable chemicals and fuels. Furthermore, our work provides initial insight into how best to combine the different alleles. Such insight is of significant importance to understanding the design principles for engineering biology, and is not gained through breeding or genome shuffling alone. Overall, our work is a step in the direction of being able to engineer polygenic complex traits from first principles.

## Methods

### Yeast Media

Yeast Extract Peptone Dextrose (YPD) media was prepared as per manufacturer directions (BD Difco, 24280) and was supplemented with the appropriate drug (200 mg/L G418, 100 mg/L clonNat (WERNER BioAgents, GmbH, 5.004.000), and/or 300 mg/L hygromycin B (Invitrogen, 10687010) when selective conditions were required. Magic medium plates^59^, used in the segregant pool generation, were prepared with 2% (w/v) D-glucose (Sigma, G7528), 0.67% (w/v) Yeast Nitrogen Base w/o Amino Acids (BD Difco, 291940), 0.2% (w/v) Drop-out Mix Synthetic Minus Arg, His, Lys, w/o Yeast Nitrogen Base (US Biologicals, D9539-05C), 50 mg/L thialysine (Sigma, A2636), 50 mg/L canavanine (Sigma, C9758), 2% Bacto Agar (BD Difco, 214010). SC80 media used in phenotypic growth assays was a synthetic complete media containing 8% (w/v) D-glucose (Sigma, G7528), 0.67% (w/v) Yeast Nitrogen Base w/o Amino Acids (BD Difco, 291940), 0.2% (w/v) Drop-out Mix Complete w/o Yeast Nitrogen Base (US Biologicals, D9515), and the pH was adjusted to 5.5 with KOH. For growth assays performed in bioreactor vessels, Antifoam 204 (Sigma, A8311) was added to a final concentration of 0.005% (vol/vol). For growth assays performed in microplates, SC80MES media was prepared by buffering SC80 media with MES (Sigma, M5287) and adjusting to pH 5.5; for the initial phenotyping of APA5026 and APA5030 in 48-well microplates, the concentration of MES was 20 mM, but for Reciprocal Hemizygosity Analysis and screening of bulk variant replacement strains in microplates, the concentration of MES was increased to 100 mM.

Plant hydrolysate, prepared by a dilute-acid pretreatment method of *Miscanthus x giganteus*, was obtained from the National Renewable Energy Laboratory^13^. The pH of this hydrolysate was adjusted to 5.5 with potassium hydroxide (KOH); in order to reduce unnecessary dilution of the hydrolysate, solid KOH was primarily used and final adjustments were made with a 15 M KOH solution. Precipitate was removed by two rounds of centrifugation at 8000 RPM in a Beckman Coulter JLA8.1000 rotor for 25 minutes at 23°C, and the clarified hydrolysate was sterilized by vacuum filtration through a 0.22 µm PES membrane (Corning, 431098). We define this filtered hydrolysate as 100% hydrolysate.

### Yeast strains

The *S. cerevisiae* strains used in this study are listed in **Table S2**. An S288C strain (ATCC^®^ 204508™, APA2574) was purchased from ATCC. First, we induced mating type switching in APA2574, allowed selfing to occur, and isolated a diploid version of S288C (APA2608). We then sporulated APA2608 and isolated APA2656, APA2657, APA2658, and APA2659 from a single tetrad. The strain referred to as S288C in the results of this study is APA2657. One-step gene replacement was used to delete one copy of the *HO* gene in APA2608, creating APA5005, which was then sporulated to obtain a *MAT*α *hoΔ::HphMX4* strain (APA5018). To generate our S288C^mm^ strain (*MAT*α *hoΔ::HphMX4 can1Δ::prSTE2-SpHIS5 lyp1Δ his3Δ1::NatMX4*), a series of genetic crosses were performed to introduce the *can1Δ::prSTE2-SpHIS5* and *lyp1Δ* alleles from APA2507 (Y7092^60^, a gift from Charlie Boone, University of Toronto) into the S288C background. First, a *MAT***a** haploid containing the *can1Δ::prSTE2-SpHIS5* and *lyp1Δ* alleles (APA5009) was obtained from a cross between APA2507 and APA2658. APA5009 was then crossed with APA5018, and the resulting diploid was sporulated. A *MAT*α haploid containing the *hoΔ::HphMX4*, *can1Δ::prSTE2-SpHIS5*, and *lyp1Δ* alleles was isolated, in which the *his3Δ1::NatMX4* allele was subsequently made by one-step gene replacement, yielding APA5019. APA5029, APA5030, and APA5031 were isolated from single colony purification of APA5019. The wild type JAY291 and a *his3Δ::KanMX* mutant version (APA2373 and APA2512, respectively) were gifts from Yong-Su Jin (UIUC). We attempted to delete the *HO* gene in APA2512 with *NatMX4*, using one-step gene replacement to generate our JAY291^mm^ strain. We isolated APA5024, and single colony purification yielded APA5026, APA5027, and APA5028. Each of these strains is Nat^R^ but upon whole-genome sequencing, we determined that *NatMX4* is not present at the *HO* locus, and likely integrated randomly by non-homologous recombination. However, because JAY291 is heterothallic due to mutations in *HO*^32^, and our S288C^mm^ strain carried a complete deletion of *HO*, the residual *ho* allele in JAY291^mm^ was not problematic. APA5030 and APA5026 were mated to construct the hybrid diploid strain (APA5035) used to generate the segregant pool. The *HphMX4* marker at the *HO* locus in APA5035 was replaced with the *amdSYM* selectable marker using an *hoΔ::amdSYM* cassette, yielding APA5504. APA4028 is the mated product of APA2373 and APA2657.

All *NatMX4* deletion cassettes were amplified from pAG36^61^. The *amdSYM* cassette^62^ was synthesized and cloned into the EcoRV site of pUC57 by Genscript to create pUC57-amdSYM, which served as the template for amplifying the *hoΔ::amdSYM* cassette. Oligonucleotides used to create these deletion cassettes are listed in **Table S3**.

Reciprocal hemizygote strains were constructed by one-step gene replacement using an *HphMX4* selectable marker to delete the locus of interest from a wild type hybrid diploid parent strain (either APA5504 or APA4028, see below). Each *HphMX4* deletion cassette was amplified by PCR using pAG26^61^ as the template and oligonucleotides with 80 nt sequence extensions homologous to sequence flanking the locus targeted for deletion. The 80 nt sequences were selected such that they contained no variant sites. Yeast transformations were performed as described below. Transformants were screened for proper deletion by PCR amplification across the intended deletion site junctions. We Sanger sequenced the PCR products to determine whether the *HphMX4* deletion cassette was linked to the S288C or JAY291 alleles at known variant sites, thereby revealing whether the S288C or JAY291 allele was deleted in that transformant. All oligonucleotides used to construct and confirm reciprocal hemizygote strains are provided in **Table S3**.

APA5504 served as the wild type hybrid diploid parent strain in construction of reciprocal hemizygotes for the QTLs 04a and 04b. However, when using APA5504, we isolated many transformants with an unintended *HphMX4*-replacement of another MX4 cassette, rather than a deletion of the target locus (unpublished observations). We therefore used the marker-free APA4028 strain as the wild type hybrid diploid parent strain for all other reciprocal hemizygote strains. We also decreased the deletion size and began deleting neighboring regions in parallel, rather than performing a binary search scheme as previously described for bRHA^63^

Bulk variant replacements were performed with a linear PCR fragment (donor DNA) cassette in homology directed repair of chromosomal double strand breaks induced by Cas9. The donor DNA, which carries the desired genetic variants to be integrated into the genome, was PCR amplified from the APA2373 (JAY291) genome. Protospacer sequences used to target Cas9 were chosen and cloned into the sgRNA/Cas9 plasmids as described below. The donor DNA and sgRNA/Cas9 plasmids were co-transformed into yeast as described below. Transformants were screened for successful bulk variant replacements using PCR to amplify the bulk variant replacement regions and Sanger sequencing of the PCR products to genotype each variant site within the region. All oligonucleotides used to construct and confirm bulk variant replacement strains are provided in **Table S3**. The sgRNA/Cas9 plasmids used in bulk variant replacement strain construction are provided in **Table S4**.

### Yeast transformations

Yeast was transformed with an *HphMX4*-marked deletion cassette for reciprocal hemizygote strain construction, or co-transformed with a donor DNA cassette and a Cas9/sgRNA plasmid for bulk variant replacements. Our transformation procedure was adapted from Pan and colleagues^59^. Yeast were grown overnight to saturation in 10 ml of YPD at 30°C, diluted to 0.25-0.3 OD_600_ in fresh YPD media, and cultured at 30°C to mid-log phase (0.6-0.7 OD_600_). Cells were then harvested by centrifugation in a tabletop centrifuge, and the cell pellet was washed twice in 5 ml of 0.1 M LiOAc. The cell pellet was resuspended in 0.1 to 1 ml of 0.1 M LiOAc, depending on the desired number of transformations; one to ten transformations were performed using these cells. A 0.1 ml volume of the cell suspension was incubated for 5-10 minutes at room temperature with 1 to 5 µg of a linear DNA cassette at 0.1 to 1 µg/µl, and for bulk variant replacement strains, 1 µl of a Cas9-sgRNA plasmid at ~0.5 µg/µl was included. A 500 µl volume of transformation mix (40% PEG_3350_, 0.1 M LiOAc, 0.1 µg/µl sheared and heat denatured salmon sperm DNA (Ambion, AM9680), 9% DMSO) was added to the cell/DNA mixture and incubated for 30-45 minutes at 30°C with shaking. Heat shock was performed at 42°C for 15 minutes. Cells were pelleted by centrifugation at 3000 RPM for 5 minutes in a microcentrifuge, resuspended in 200 µl of YPD, recovered for 1.5-4 hours at 30°C with shaking at 200 RPM, and plated on the appropriate selective media; the CaCl_2_ step described by Pan and colleagues^59^ was typically skipped.

### Whole genome sequencing and read mapping

All yeast genomic DNA extractions were performed as described previously^64^. Whole genome sequencing was performed by the UC Davis Genome Center (Davis, CA) using the Illumina HiSeq2000 platform to produce 100 bp paired end reads. The whole genome sequence data from this publication have been submitted to the NCBI Sequence Read Archive and assigned the identifier PRJNA322647. Reads were mapped to the S288C reference genome (v. R64-1-1, release date Feb 3, 2011) as described previously^65^, except that updated versions of Picard (v. 1.103(1598)) and the GATK (2.7-4-g6f46d11)^66^ were used in this work.

### Variant detection

Variant detection and filtering were performed as described previously^65^, except that both per-sample and multi-sample variant detection were used. To construct the set of ~46,000 segregating sites between our two parent strains that we used as markers in our QTL analysis (**File S2**), we created a variant call set for each of our parent strains, relative to our S288C reference genome (**File S1**), and then took the discordant SNP sites between these two strains. Per-sample variant detection and filtering was performed for three isolates of JAY291^mm^ (APA5019, APA5026, and APA5028), each sequenced separately. To further help reduce false positive variant sites, we took the intersection of these variant call sets (i.e. only the variants that appeared in all three filtered call sets). Our S288C^mm^ call set was generated with multi-sample variant detection using three isolates of S288C^mm^ (APA5024, APA5030, and APA5031), each sequenced separately, and we did not filter this S288C^mm^ call set. Finally, we selected all SNP sites in our JAY291^mm^ call set that were absent from the S288C^mm^ call set. Mean sequencing coverage for APA5019, APA5024, APA5026, APA5028, APA5030, and APA5031 was 484, 548, 164, 168, 178, and 176, respectively.

### Bioreactor fermentations

Anaerobic fermentations were performed using a DASGIP Parallel Bioreactor System with 1 L Bioblock Stirrer vessels (Eppendorf AG Bioprocess Center, Juelich, Germany). Temperature, pH, dissolved oxygen, and impeller speed were continuously monitored and maintained at 30°C, 5.5, 0% saturation, and 400 RPM, respectively. Vessels were sparged with N_2_ gas to create an anaerobic environment (6-12 sL/h based on a DO cascade in both **Figure S1**, and the QTL experiment with segregant pool. Constant 12 sL/h was used in characterization of engineered strains). Culture density was also continuously monitored during fermentations using an optical density monitoring system (Fogale Nanotech, Nîmes, France), and this information was communicated to the DASware software via the Open Platform Communications (OPC) protocol. For continuous-culture fermentations, the DASGIP system was operated as a turbidostat; a customized script, obtained from Eppendorf DASGIP (Eppendorf AG Bioprocess Center, Juelich, Germany), controlled addition of fresh media into the bioreactor vessels in response to increasing OD values, and the removal of culture from the bioreactor vessels in response to increasing volume (https://www.eppendorf.com/uploads/media/Application-Note_No_298_DASware-analyze_Automated-Bioreactor-Sampling.pdf). This control system allowed a defined OD set point to be maintained throughout the course of the experiment (**Figure S2**). One-sided pH control with KOH was used in continuous culture fermentations. Unless otherwise noted, two-sided pH control using KOH and H_2_SO_4_ was employed in batch fermentations characterizing the engineered strains.

### Generation and growth of the segregant pool

The hybrid diploid strain, APA5035, was grown overnight in YPD at 30°C, diluted back to OD_600_ = 0.225 in fresh YPD, and grown at 30°C to ~0.6 OD_600_. Cells were harvested by centrifugation at 6000 RPM in a Beckman Coulter JLA-8.1000 rotor for 10 minutes at 23°C, washed twice in sterile ddH_2_O, resuspended at OD_600_ ~0.6 in 1% potassium acetate, and incubated for 22°C with shaking at 250 RPM to induce sporulation. Sporulation was allowed to proceed for 13 days, at which time greater than 80% of the culture had sporulated. The *MAT***a** haploids were isolated from the sporulation mixture essentially as described previously^17^, except that Zymolyase (Zymo Research, E1005) was prepared in 0.5X PBS pH 7.4, 50% Glycerol and used in place of β-glucuronidase. Briefly, sporulated and unsporulated yeast were collected by centrifugation, incubated with Zymolyase at 30°C and vortexed well with acid washed glass beads (Sigma, G8772) to liberate ascospores and spheroplast unsporulated diploids. The mixture was then diluted 20-fold in sterile ddH_2_O, and plated on magic medium. Germinated yeast was scraped from magic medium plates, suspended in 15% glycerol, pooled, aliquoted in convenient volumes and frozen at −80°C.

To enrich for segregants with superior fitness in hydrolysate, we subjected the segregant pool to continuous-culture anaerobic fermentation (see above) in SC80 media containing 30% (vol/vol) dilute-acid treated *Miscanthus x giganteus* plant hydrolysate. As a control, a fermentation lacking hydrolysate was performed in parallel. To this end, an aliquot of the segregant pool was recovered from storage at −80°C by aerobic cultivation in SC80 media at 30°C for 3.5 hours. The culture was split, cells were harvested and used to inoculate bioreactor vessels containing either SC80 or SC80 + 30% (vol/vol) hydrolysate media to OD_600_ = 0.2 (1 cm path length). The T_initial_ samples were collected immediately after inoculation. The culture densities were allowed to increase to OD_880_ = 5 (0.5 cm path length), at which point, turbidostat operation commenced, which held the culture densities and volumes constant for the remainder of the fermentations. Both the experimental and control fermentations were cultured for 75 population doublings, which was determined from the amount of fresh media added and the culture volumes. After 75 population doublings, the fermentations were stopped and T_final_ culture samples were collected.

### SNP allele frequency calculations

We used MULTIPOOL^31^ to calculate the observed and inferred SNP allele frequencies that appear in all plots presented in this work. We isolated genomic DNA from the samples collected in our pool selection experiment, performed whole genome sequencing, and mapped the reads as described above. Mean sequencing coverage for the pre-selection samples from the control and experimental pools was 182 and 179, respectively. Mean sequencing coverage for the post-selection samples from the control and experimental pools was 197 and 231, respectively. To prepare our data for analysis with MULTIPOOL, we created vcf files from our mapped reads using samtools mpileup with the -u, -D, -S, and -B options, and bcftools view with the -c and and -g options (samtools version 0.1.19-44428cd). We then extracted S288C and JAY291 SNP allele counts for our ~46,000 segregating sites from the vcf files using the DP4 tag and the GATK VariantsToTable walker. From these SNP allele counts, we calculated the observed and inferred SNP allele frequencies using MULTIPOOL (version 0.10.1) with the following parameters: -n 1000, -c 3300, -r 100.

### QTL Detection

We used the method of Parts and colleagues^16^ (iQTL), to identify large chromosomal regions containing QTLs, and subsequently analyzed those regions with MULTIPOOL to obtain high-resolution predictions for the causative QTL region. Our motivation for the sequential implementation of these two QTL detection methods was to take advantage of MULTIPOOL’s highly sophisticated approach for mapping QTLs while overcoming its limitation of being able to call only one QTL per analyzed region (e.g. per chromosome).

To prepare our data for iQTL analysis, we generated pileup files for our set of ~46,000 segregating variant sites from the mapped reads using samtools pileup (version 0.1.12a (r862)) with the -c, and -s options, and the -m and -N parameters set to 0x704 and 2, respectively. We also included the -B flag to prevent the original base quality scores from being overwritten with the calculated base alignment quality score. From these pileup files, we used Python scripts generously provided by Leopold Parts (University of Toronto) to calculate the observed and inferred SNP allele frequencies at the segregating sites according to the iQTL statistical model, to call QTLs, and for each QTL, to identify all associated loci that experienced a significant change in SNP allele frequency (i.e. define the region representing the base of a QTL peak). We set the parameters in the calculation of the inferred SNP allele frequencies to account for our use of the haploid progeny from the first filial (F1) generation in our QTL mapping population, and to increase the cutoff for probability of linkage to 0.95. We also adjusted the SNP allele frequency change cutoff in accordance with our data; this cutoff is one of the two criteria used in calling QTLs by Parts and colleagues^16^, and our cutoff value was chosen such that the probability of observing an SNP allele frequency change of at least equal magnitude in our control experiment would be less than 0.025.

For each QTL-containing region identified with the iQTL method, the SNP allele counts we previously collected for the calculation of SNP allele frequencies with MULTIPOOL were filtered to contain only the variant sites present within the identified region. The filtered SNP allele counts for the post-selection samples from the control and experimental pools were simultaneously analyzed with MULTIPOOL (-m option set to contrast, all other parameters as before) to produce the high-resolution QTL map.

### Cloning of protospacers into sgRNA/Cas9 vectors

Single-guide sgRNA/Cas9 plasmids were constructed by cloning protospacer sequences into pCas9 (a generous gift from Jamie Cate, UCB) using restriction free cloning as described previously^65,67^. Dual-guide sgRNA/Cas9 plasmids were constructed by Golden Gate Assembly according to Lee and colleagues^68^. Plasmid pML1414 (a generous gift from John E. Dueber, UCB) served as the Type 234 sgRNA dropout vector. This Type 234 sgRNA dropout vector differs from the one described in Lee and colleagues^68^ in that expression of sgRNAs are regulated by the SNR52 promoter and SUP4 terminator described by DiCarlo and colleagues^69^, the GFP expression cassette is flanked by BbsI sites, and it requires an upstream overhang of 5’GATC in the annealed oligonucleotides. Oligonucleotides used in plasmid constructions are listed in **Table S3**. The final sgRNA/Cas9 plasmids constructed and used to generate bulk variant replacement strain are provided in **Table S4**.

To select the protospacer sequences, we generated lists of all potential Cas9 targeting sequences (the 23 nt sequences corresponding to each PAM site (NGG) and the immediate upstream 20 nt protospacer sequence) within the intended bulk variant replacement region that contained at least one SNP within the 23 nt targeting sequence, not including SNPs at the N position in the PAM. For dual-guide sgRNA/Cas9 plasmids, which were used in the QTL 10a and QTL 12c swaps, two protospacer sequences were selected to maximize the number of variant sites in the bulk variant replacement region flanked by the two resulting Cas9 double-strand breaks sites. If more than one protospacer sequence satisfied this criterion, then we selected the protospacer sequence corresponding to the 23 nt Cas9 targeting sequence with a SNP (excluding those at the N position of the PAM) closest to the 3’ end of the Cas9 targeting sequence.

### Microplate reader growth assays for hydrolysate tolerance screening

Each RHA and bulk variant replacement strain in each condition was assayed in at least three independent replicate experiments. For some bulk variant replacement genotypes, data was collected from multiple independently constructed strains. In each experiment, a saturated SC80MES culture of yeast, grown aerobically at 30°C was prepared for each strain. Cells were harvested by centrifugation in a microcentrifuge, resuspended to OD_600_ = 1 in 5X SC80MES media and transferred to an anaerobic chamber (Coy Laboratory Products, Inc). The cell suspension was diluted 5-fold into wells of a microplate containing a dilution series of hydrolysate. Microplates were covered with a gas-permeable plate sealing membrane: “Breathe-Easy” sealing membranes (USA Scientific, 9123-6100) for reciprocal hemizygosity experiments or Mylar Plate Sealer sealing membranes (MP Biomedicals, 76-402-05) for bulk variant replacement experiments, and then placed into a plate reader within the anaerobic chamber. Unless otherwise noted, Tecan Sunrise^TM^ microplate readers were used to measure the optical density of the culture at 595 nm or 600 nm every 15 minutes with shaking, using the “Normal” speed setting, performed between measurements.

### Growth curve data analysis

All microplate growth data, and code used for growth data analysis is available in the **File S3**. Data collected from all strains of the same genotype were used in all calculations. Raw optical density measurements from microplate reader growth assays were background-corrected by subtracting the within-plate mean optical density values of wells containing media only. No path length correction was performed since the same culture volume (150 µl) was used in all wells of all microplate growth plate assays. Background-corrected optical density values were used in plots of OD vs. Time. We occasionally observed in some trials (i.e. wells), massive evaporation accompanied by cells dispersing to the edges of the well and causing an artificial, complete decay of the measured OD. This phenomenon usually followed a period of significant growth, and likely resulted from a compromised plate seal for that particular well caused by the CO_2_ gas production rate during fermentation exceeding the permissible gas exchange rate for the sealing membrane. OD measurements from periods during which this artificial decay occurred were excluded from plots of OD vs. Time.

Two methods were used to determine the maximum culture densities used in the comparisons of hydrolysate tolerances between genotypes. The first method used cubic splines to smooth the background-corrected growth data (**File S3**) for each trial of each strain in each condition. Then the maximum OD value for each trial was obtained from the smoothed data, and the mean maximum culture density (max OD) was calculated for each genotype in each condition.

In addition to comparing maximum culture densities, we also compared changes in lag phase and maximum growth rates. We generated (four-parameter) logistic growth curve models (**File S3**) describing these parameters for each trial, and obtained parameter estimates from the fitted models. In order to obtain model fits that accurately described max culture density, lag, and max growth rates, we reduced the influence of the artificial decrease in measured OD during stationary phase on the model fits by determining the time at which the maximum OD value was observed in the smoothed data, and using the background-corrected OD value from that time point for all remaining time points in the trial. This method is similar to a previous method for removing artificial negative slopes in growth curve data^70,71^, but has greater data conservation/fidelity. The logistic growth curve model was fit to each trial by nonlinear least squares regression using the nls() function in R (**File S3**). The parameter estimates for each trial were obtained from the fitted models, and the means were calculated for each strain in each condition.

The smoothing-splines method could be implemented for every strain in every condition, whereas it was difficult to obtain a logistic growth curve model fit in some instances, particularly when strains were completely unable to grow in a given hydrolysate concentration. We therefore present the maximum OD as calculated by the smoothing-splines method in any case where the genotypes being compared included one or more other strains completely unable to grow in the given condition. It is important to note that regardless of which method is used, the same conclusions can be reached for all genotype comparisons.

To calculate the mean relative maximum culture densities (relative max OD), we first obtained the maximum culture densities, using the smoothing-splines method, for each trial in the microplate growth assay. We then normalized each trial to the S288C trials performed within the same microplate, and the mean relative maximum culture densities (relative max OD) were calculated from the normalized values.

### Characterizing hydrolysate tolerance of engineered strains in a bioreactor

Yeast was cultured aerobically overnight to saturation in SC80MES at 30°C. Saturated cultures were diluted to OD_600_ = 0.2 in fresh SC80 and grown aerobically at 30°C for 30 hours. Cells were harvested and used to inoculate SC80 supplemented with 40% (vol/vol) hydrolysate in the bioreactor at OD_600_ = 0.25 (1 cm path length) for an anaerobic batch fermentation. Culture density was continuously monitored and measurements were logged at least every two minutes (as described above). Cell-free samples were collected every 8 hours for analysis of glucose utilization and ethanol production. Sample collection was performed using a Flownamics autosampling system comprised of a SegFlow4800, Flowfrac400 fraction collector, and FISP probes. Samples were filtered and analyzed using a 1200 series high performance liquid chromatography system (Agilent Technologies, Santa Clara, CA, USA) consisting of an autosampler with tray cooling, binary pump, degasser, thermostated column compartment and refractive index detector (RI) connected. The supernatant was injected onto a 300 mm × 7.8 mm (length × inner diameter) Aminex HPX-87H (Bio-Rad, Richmond, CA, USA) column with 9 µm particle size, 8% cross-linkage, equipped with a 30 × 4.6 mm micro-guard Cation H guard column cartridge (Bio-Rad, Richmond, CA). Compounds were eluted at 50 °C at a flow rate of 0.6 mL/minute using a mobile phase of 5 mM sulfuric acid. Integrated peak areas of glucose and ethanol were compared to an external calibration.

## Acknowledgments

We thank Leopold Parts for providing code for iQTL analysis, and for helpful discussion during the early stages of this work. This work was supported by the Energy Biosciences Institute Grants OO7G02 and OO1605 (A.P.A).

## Author Contributions

M.J.M., J.M.S., and A.P.A. conceived and designed the study; M.J.M., L.S., A.L.M., D.P.,

J.M.S., T.D.M., and S.B. performed the experiments; M.J.M., J.M.S., A.P.A., and S.B. analyzed the data and wrote the manuscript.

## Footnotes

† In this work, we do not limit our use of the term allele to a protein or RNA coding gene, but rather to refer to an allelic form of a genomic locus. The allelic locus might be as small as a single nucleotide position such as a SNP site, or describe a larger region such as a defined QTL, possibly containing multiple genes.

